# MAGqual: A standalone pipeline to assess the quality of metagenome-assembled genomes

**DOI:** 10.1101/2023.12.13.571510

**Authors:** Annabel Cansdale, James P.J. Chong

## Abstract

Metagenomics, the whole genome sequencing of microbial communities, has provided insight into complex ecosystems. It has facilitated the discovery of novel microorganisms, explained community interactions, and found applications in various fields. Advances in high-throughput and third-generation sequencing technologies have further fuelled its popularity. Nevertheless, managing the vast data produced and addressing variable dataset quality remain ongoing challenges. Another challenge arises from the number of assembly and binning strategies used across studies. Comparing datasets and analysis tools is complex as it requires a measure of metagenome quality. The inherent limitations of metagenomic sequencing, which often involves sequencing complex communities means community members are challenging to interrogate with traditional culturing methods leading to many lacking reference sequences.

The MIMAG standards (Bowers *et al*., 2017) aim to provide a method to assess metagenome quality for comparison but have not been widely adopted. To bridge this gap, the MAGqual pipeline outlined here offers an accessible way to evaluate metagenome quality and generate metadata on a large scale. MAGqual is built in Snakemake to ensure readability and scalability and its open-source nature promotes accessibility, community development, and ease of updates. Here, we introduce the pipeline MAGqual (metagenome-assembled genome qualifier) and demonstrate its effectiveness at determining metagenomic dataset quality when compared to the MIMAG standards. MAGqual is built in Snakemake, R, and Python and is available under the MIT License on GitHub at https://github.com/ac1513/MAGqual.

## Introduction

Metagenomics, the analysis of whole genomes of microbial communities directly from environmental samples, has proved to be a revolutionary tool in microbiology. With applications in environmental, medical and biotechnology arenas, metagenomics has resulted in the discovery of many interesting species and even whole phyla that had remained uncharacterised because they aren’t easily manipulated in the lab or are unculturable (Pelletier *et al*., 2008; Zaremba-Niedzwiedzka *et al*., 2017). This has led to the elucidation of the real dynamics of more complex microbial communities (Van Goethem *et al*., 2021).

Metagenomic sequencing of environmental samples has become increasingly popular in recent years, mainly due to the development of next-generation sequencing. Sequencing technologies have become higher throughput and lower in cost, which makes the sequencing of entire microbial communities more feasible (Albertsen, 2023). Due to the advantage metagenomic sequencing offers by removing the need for isolation and amplification of organisms, many mixed microbial communities once known as the “uncultured microbial majority” or “microbial dark matter” (Rappé and Giovannoni, 2003; Filée *et al*., 2005) that were previously challenging to characterise (Nayfach *et al*., 2021), have been targeted by metagenomic sequencing. A result of metagenomics targeting little-characterised communities is that many organisms found in metagenomics studies have either never been seen before or have previously been uncultured so lack available reference genomes (Nayfach *et al*., 2021).

A typical metagenomics analysis pipeline would be as follows: raw reads from shotgun DNA sequencing of a microbial community would undergo quality control before being assembled using appropriate assembly software (including MetaSPAdes, MEGAHIT, IDBA-UD for short-read metagenomics and metaFlye, Canu for long-read metagenomics (Peng *et al*., 2012; Li *et al*., 2015; Koren *et al*., 2017; Nurk *et al*., 2017; Kolmogorov *et al*., 2020)), then the metagenomic assembly would be binned using a variety of approaches to group the contigs associated with different organisms in the sequenced community into metagenome-assembled genomes (MAGs) (Pérez-Cobas, Gomez-Valero and Buchrieser, 2020).

Metagenomic-specific software employs many different assembly and binning strategies (Pérez-Cobas, Gomez-Valero and Buchrieser, 2020) as metagenomic studies have different challenges than single-organism genomic studies. A mixed community with organisms at different abundances makes both the assembly and binning of a metagenome challenging (Nurk *et al*., 2017; Pasolli *et al*., 2019; Pérez-Cobas, Gomez-Valero and Buchrieser, 2020). The risk of contamination of MAGs from closely related organisms is an additional challenge (Pérez-Cobas, Gomez-Valero and Buchrieser, 2020). Therefore, it is important to have a method of determining overall metagenome and MAG quality.

The lack of reference genomes available for many organisms identified through metagenomics becomes an issue when comparing metagenome analysis methods and software due to a lack of ground truth. Benchmarking and quality assessment tools exist for metagenomic studies, such as AMBER and MetaQUAST (Mikheenko, Saveliev and Gurevich, 2016; Meyer *et al*., 2018), however, these require the organisms present in the dataset to be known and have appropriate reference genomes.

Determining MAG quality is important to indicate the quality of the initial analysis and highlight which MAGs are worthy of further investigation or deposition onto online databases. The Minimum Information about a Metagenome-Assembled Genome (MIMAG) (Bowers *et al*., 2017) is a standard developed by the Genomics Standards Consortium (GSC) which outlines a framework for the classification of MAG quality (into either high-quality draft, medium-quality draft or low-quality draft) and recommends the reporting of specific metadata for each MAG. While this framework aids in the reproducibility of metagenomic studies it has not yet received universal uptake.

Within the MIMAG standards, three criteria are used to determine overall MAG quality: genome completeness, contamination and assembly quality. When taxonomy is known and a reference genome is available for a MAG, these metrics are easier to determine. However, identifying and pairwise-alignment of MAGs to a reference is a manual and computationally intensive process so is not an appropriate method for a large number of MAGs (Pérez-Cobas, Gomez-Valero and Buchrieser, 2020). Further to this, as metagenomic sequencing is often used to shotgun sequence environmental samples, it captures many organisms that are a part of the microbial dark matter and as a result, do not have closely related reference genomes (Pasolli *et al*., 2019).

Due to the lack of a “ground truth” (i.e. a closely related reference strain) for many communities that are investigated using metagenomic sequencing and the computational power required to determine closely related organisms at scale, it is necessary to take a reference-free or *de novo* approach to determine the success of both metagenomic sequencing and binning (Bowers *et al*., 2017). One such approach determines the completeness and contamination of a genome (or in this case a metagenome-assembled genome) using marker genes, as exemplified by the popular software CheckM (Parks *et al*., 2015). For many organisms identified through metagenomics, determining assembly quality is challenging as there is not a defined sequence to compare the MAG back to. For MAGs, determining assembly quality is suggested to be determined by the presence and completeness of encoded rRNA and tRNA genes within the metagenome bin (Bowers *et al*., 2017).

Due to the abundance of metagenomic software available, community adoption of standards like MIMAG is important to increase reproducibility and reliability within and between datasets, however, currently, the MIMAG standards remain under-utilised by many studies. The advances and increasing throughput of metagenomic sequencing have resulted in the generation of hundreds to thousands of MAGs per metagenome (Parks *et al*., 2017; Pasolli *et al*., 2019). Parsing the information required to determine the quality of these bins and isolating the higher-quality MAGs worthy of further analysis is a challenge. It was deemed useful to produce an analysis pipeline to automate this at scale.

Here we introduce MAGqual (**M**etagenome-**A**ssembled **G**enome **Qual**ity), a pipeline implemented in Snakemake v7.30.1 (Mölder *et al*., 2021). MAGqual enables the user to pass in MAGs generated by metagenomic binning software and quickly assess the quality of these bins according to the MIMAG standards. These bins are analysed to determine completeness and contamination (using CheckM v1.0.13 (Parks *et al*., 2015)) and the number of rRNA and tRNA genes (using Bakta v1.7.0 (Schwengers *et al*., 2021)) that each bin encodes. This information is used by bespoke code to determine the quality of each bin, in line with the MIMAG standards (with an additional “near complete” category) and produces figures and a report that outlines the quality and other metrics of the input MAGs.

MAGqual enables users to automate the assignment of quality to their metagenome bins and quickly determine the success of their metagenomic analysis and will hopefully improve the uptake of MIMAG standards across the metagenomics community and provide an easy way to benchmark new metagenomic binning software or analysis methods. Its open-access nature and simple Snakemake pipeline will enable timely updates as the metagenomic field moves forward. MAGqual is available from https://github.com/ac1513/MAGqual under an MIT license.

## Methods

### MAGqual pipelines

The MAGqual pipeline is built in Snakemake (v.7.30.1) (Mölder *et al*., 2021). Snakemake is a popular workflow management tool based on Python which enables a human-readable plug-and-play strategy for analysis pipeline design. This method of design results in a pipeline that is easier to understand, adapt, and maintain. Furthermore, Snakemake integrates easily into high-performance computing clusters, making workflows highly scalable on many systems - which is key as datasets increase in size.

The MAGqual pipeline requires minimal installation, only Miniconda and Snakemake are required to be in the user’s environment as the installation of all other software is handled by the Snakemake pipeline with Conda environments. MAGqual handles the installation of databases required by Bakta and CheckM. The light version of the Bakta database is downloaded to maximise speed and minimise storage space required, however, MAGqual allows the specification of a local Bakta and/or CheckM database if required. Two file types are required as input: first, the metagenomic bins (MAGs) in FASTA format (with the file extension: fasta, fna, or fa), and second the metagenomic assembly (in FASTA format) used to generate the metagenomic bins.

To remain accessible to those unfamiliar with Snakemake pipelines, MAGqual can be run using a Python wrapper, with the basic command:

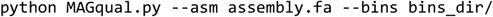

Full pipeline functionality can be achieved using this Python wrapper, which prevents users from needing to edit configuration files. See Table 1 for the full command line options available to the user and their defaults.

**Table 1:** MAGqual command line options and their usage.

| Command line option | Usage |
| --- | --- |
| -a / --asm | **Required:** the location of the assembly in FASTA format used to generate the metagenome bins |
| -b / --bins | **Required:** the location of the directory containing all the metagenome bins for quality assignation |
| -p / --prefix | The prefix for the job. Default is MAGqual_YYYYMMDD |
| -j / --jobs | Links to the Snakemake flag -j, the number of cores to use or if using the cluster option, the number of jobs to run at once. Default is 1. |
| --cluster | Optional: The type of cluster to run MAGqual on a HPC system (available options: slurm), not to be used if running MAGqual locally. |
| --checkmdb | Optional: The location of a local install of the CheckM database. |
| --baktadb | Optional: The location of a local install of the Bakta database. Note: must be v.5 or above. |
| -h / --help | Show help message. |

**Table 2:** MAGqual quality evaluation categories and the completeness, contamination and rRNA/tRNA requirements.

| Quality | Completeness | Contamination | rRNA/tRNA required |
| --- | --- | --- | --- |
| High | > 90% | $\leq 5\%$ | $\geq 18$ tRNA and 23S, 16S, and 5S rRNA |
| Near complete | > 90% | $\leq 5\%$ | None |
| Medium | $\geq 50\%$ | $\leq 10\%$ | None |
| Low | < 50% | $\leq 10\%$ | None |
| Failed | - | $\geq 10\%$ | None |

MAGqual also retains full Snakemake functionality and can easily be run using the Snakemake architecture. The basic command for this is, snakemake `--use-conda -j 1` and the user is required to edit the `config/config.yaml` file to specify the location of the input files and has the option to further edit the command and the `config/cluster.json` file to add configuration options to run MAGqual on an HPC cluster. This is a useful option for those more familiar with Snakemake pipelines, as the pipeline can be further modified to run better on different infrastructures however this is not necessarily appropriate for every user.

### MAG completeness and contamination

While MIMAG introduces metagenomic standards it does not recommend a specific “industry-standard” method of calculating them. Both the completeness and contamination values vary depending on what set of initial single-copy marker genes is used, so the method used requires reporting. Here we use the software CheckM which in the years since the publication of the standards has become the de facto software used for these calculations (Yang *et al*., 2021).

### Assembly Quality

In line with the MIMAG standards, the presence and completeness of tRNA and rRNA ribosomal genes are also determined. This is suggested to be a method of assembly quality determination. Here, the MAG is run through Bakta for annotation and rapidly identifies rRNA and tRNA ribosomal genes. Previously, Prokka has been the software of choice for fast microbial annotation (Seemann, 2014) however Bakta improves the annotation of CDS compared to Prokka and remains actively supported so it was chosen for use in this pipeline (Schwengers *et al*., 2021). To be classified as a high-quality draft MAG, a MAG must encode tRNAs for at least 18 of the 20 possible amino acids and the 5S, 16S and 23S rRNA genes (Bowers *et al*., 2017).

### Determining MAG quality

Using the results from CheckM and Bakta, MAGqual calculates overall MAG quality using Python. Along with the high, medium, and low-quality standards introduced in the MIMAG standards (Bowers *et al*., 2017), we include the “near complete” quality draft MAG category introduced by (Almeida *et al*., 2019). We define this “near complete” quality (NCQ) draft MAG as >90% complete, <5% contaminated but not encoding the necessary tRNA and rRNA to be classified as a highly complete draft MAG. This was determined to be an important addition to MAGqual due to documented problems around the assembly and annotation of rRNA/tRNA sequences (Yang *et al*., 2021), especially with metagenomes generated from short-read sequencing (Parks *et al*., 2017; Singleton *et al*., 2021). As uncultured and previously undetermined organisms make up a significant proportion of metagenomic communities this enables some flexibility for any CDS annotation issues.

Once the quality of the MAGs has been calculated a figure showing a breakdown of size, completeness, and contamination scores for all bins, as well as a quality category is generated to provide the user with simple and quick evidence of the quality of their metagenome bins. A file containing the recommended metadata (see Table 3) for each of the MAGs is exported into one file to enable easy submission and analysis. An interactive HTML report is also generated to allow easy viewing of these plots and metadata. MAGqual will finally output multiple directories containing the MAGs split by overall quality category.

**Table 3:**
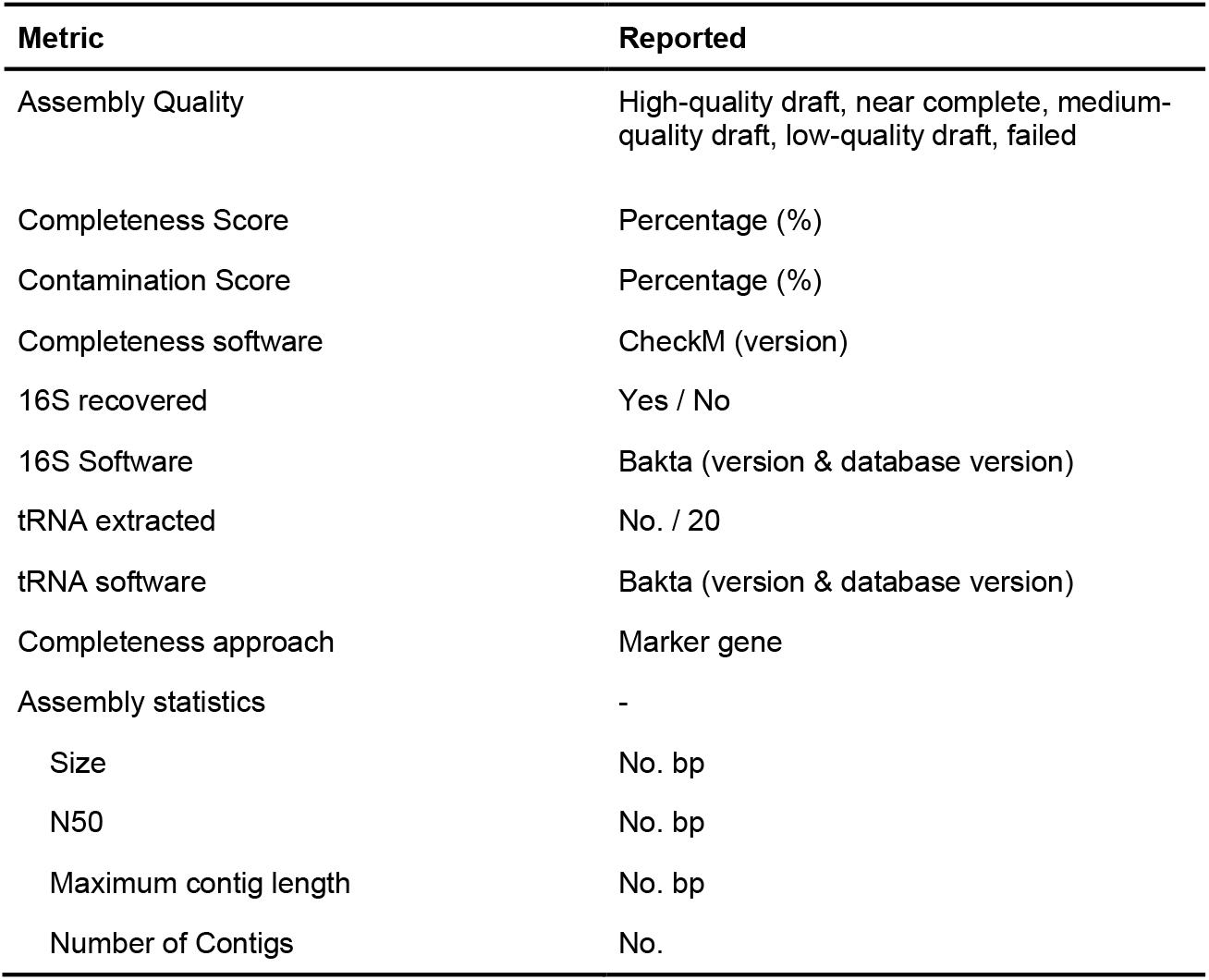
Metadata and metrics reported by the MAGqual pipeline in CSV format.

| Metric | Reported |
| --- | --- |
| Assembly Quality | High-quality draft, near complete, medium-quality draft, low-quality draft, failed |
| Completeness Score | Percentage (%) |
| Contamination Score | Percentage (%) |
| Completeness software | CheckM (version) |
| 16S recovered | Yes / No |
| 16S Software | Bakta (version & database version) |
| tRNA extracted | No. / 20 |
| tRNA software | Bakta (version & database version) |
| Completeness approach | Marker gene |
| Assembly statistics | - |
| Size | No. bp |
| N50 | No. bp |
| Maximum contig length | No. bp |
| Number of Contigs | No. |

#### Reported metadata

Alongside determining MAG quality, MAGqual also generates metadata recommended by the MIMAG standards. The metadata categories can be seen in Table 3, MAGqual produces a CSV file with a line of corresponding metadata for each MAG run through the pipeline.

#### MAGqual Report

Along with a metadata table, MAGqual generates an interactive HTML report generated using Python, RMarkdown and Plotly. This report produces numerous figures including the bases and contigs binned, completeness and contamination, N50 length, total length and tRNA completeness along with tables for MAG quality and metadata. This report is generated from all available MAGqual runs in the same directory, enabling a quick comparison of binning results between MAGqual runs.

#### Running the pipeline

To determine the runtime of the pipeline on minimal architecture, an 8-core 32GB Linux instance (Canonical Ubuntu 22.04) on Oracle Cloud was used. Snakemake (v7.30.1) and Conda (v.23.5.2) were installed into this nascent environment and MAGqual handled the installation and database downloads required.

Due to the requirement for over 40GB of memory, CheckM was run for this test with the -- reduced_tree option which lowers the memory required to 16GB (see https://github.com/Ecogenomics/CheckM/wiki/Installation#system-requirements).

To reflect actual runtime each benchmark was run from a clean environment, therefore the Conda environments had to be remade and the databases re-downloaded to include the time required for these into each run.

To validate the MAGqual pipeline we generated a dataset of 10, 100, 500 and 1,000 MAGs from Parks *et al*. (2017). This study was chosen as these MAGs were previously assigned a quality with completeness and contamination scores. However, this paper predates the publication of the MIMAG standards so no high-quality MAGs were defined.

#### Comparison of binning tools

A small metagenomics dataset from a gut microbiome was procured from ENA project PRJEB44880 (Shahi *et al*., 2022) corresponding to a Nanopore metagenome assembly polished with Illumina short-reads and seven samples of short-read Illumina raw sequencing data. Three popular metagenomic binning tools were chosen for comparison, CONCOCT (v1.1.0), Metabat2 (v2.12.1) and Binsanity (v0.5.4) (Alneberg *et al*., 2014; Graham, Heidelberg and Tully, 2017; Kang *et al*., 2019) all three tools use abundance information for binning which was generated using BWA (v0.7.17) (Li, 2013) to map the Illumina short reads back to the assembly generating a BAM file for each short-read sample.

These BAM files were then passed to each binning software using the minimum contig length of 1000 bp - apart from CONCOCT which requires contigs <10 kb where the assembly had to be split and the raw reads re-mapped to each split assembly. The MAGs produced by these three binners were passed through the bin refinement tools MetaWrap (v1.3.2) and DAS tool (v1.1.6) (Sieber *et al*., 2018; Uritskiy, DiRuggiero and Taylor, 2018). DAS tool was run as directed, with the flag --write_bins to produce the bins for comparison. MetaWrap was run using the bin_refinement module and the flags -c 0 (minimum completeness) -x 100 (maximum contamination) to output all bins regardless of quality.

## Results

### Running the pipeline

MAGqual is quick and easy to run, with only one command required to initiate the pipeline, install dependencies, and run each of the steps. The speed of the pipeline depends on the size of the dataset being analysed and is overall limited by the speed of the programs Bakta and CheckM.

As seen in Table 4, with the generated dataset of 10, 100, 500 and 1,000 MAGs from (Parks *et al*., 2017), as the number of MAGs increases most of the run time is assigned to Bakta, which, while only taking on average ~2.5 minutes per MAG, becomes substantial when running 1000 MAGs.

**Table 4:** Time and memory requirements for the MAGqual pipeline when run with an increasing number of MAGs (10, 100, 500 and 1000) on an 8-core 32GB RAM instance. The real-time, CPU time and maximum memory of the overall pipeline and the individual steps of Bakta. CheckM and Python scripts are outlined below. As Bakta runs a separate job for each MAG the average real-time, average CPU time and average max memory are also included.

|  |  | MAGqual overall |  | Bakta |  |  |  | CheckM |  | Scripts |  |
| --- | --- | --- | --- | --- | --- | --- | --- | --- | --- | --- | --- |
|  |  | Wall clock Time (HH:MM:SS) | CPU time (HH:MM:SS) | Total Wall clock Time (HH:MM:SS) | Total CPU Time (HH:MM:SS) | Average Wall clock time (HH:MM:SS) | Average CPU time (HH:MM:SS) | Wall Time (HH:MM:SS) | CPU time (HH:MM:SS) | Wall Time (HH:MM:SS) | CPU time (HH:MM:SS) |
| **No. MAGs** | **10** | 00:40:21 | 01:46:38 | 00:26:05 | 01:27:44 | 00:02:37 | 00:08:46 | 00:03:55 | 00:15:00 | 00:00:10 | 00:00:12 |
|  | **100** | 04:41:35 | 16:05:00 | 04:13:00 | 16:40:26 | 00:02:32 | 00:10:00 | 00:25:28 | 02:51:03 | 00:00:09 | 00:00:09 |
|  | **500** | 22:59:02 | 84:18:59 | 20:58:15 | 77:18:18 | 00:02:31 | 00:09:17 | 03:31:57 | 11:06:23 | 00:00:12 | 00:00:10 |
|  | **1000** | 45:48:35 | 164:08:03 | 41:56:34 | 95:31:20 | 00:02:31 | 00:11:44 | 04:27:48 | 29:30:18 | 00:00:15 | 00:00:11 |

While it was important to determine the minimal computational infrastructure required to run this pipeline, as the majority of metagenomic research is undertaken on HPCs it is recommended to run Bakta jobs in parallel which is possible using the flag `-j` using both the Python wrapper and using the Snakemake infrastructure.

### Validation of MAGqual

The 1000 MAGs used in the previous analysis were also used to compare the quality score assigned by Parks *et al*. (2017) (Fig. 2A) to the new MAGqual quality score (Fig 2B). The introduction of MIMAG quality scores changed the assignment of many MAGs from the original dataset, classifying 17 high-quality MAGs which were previously within the “near complete” category.

**Figure 1:**
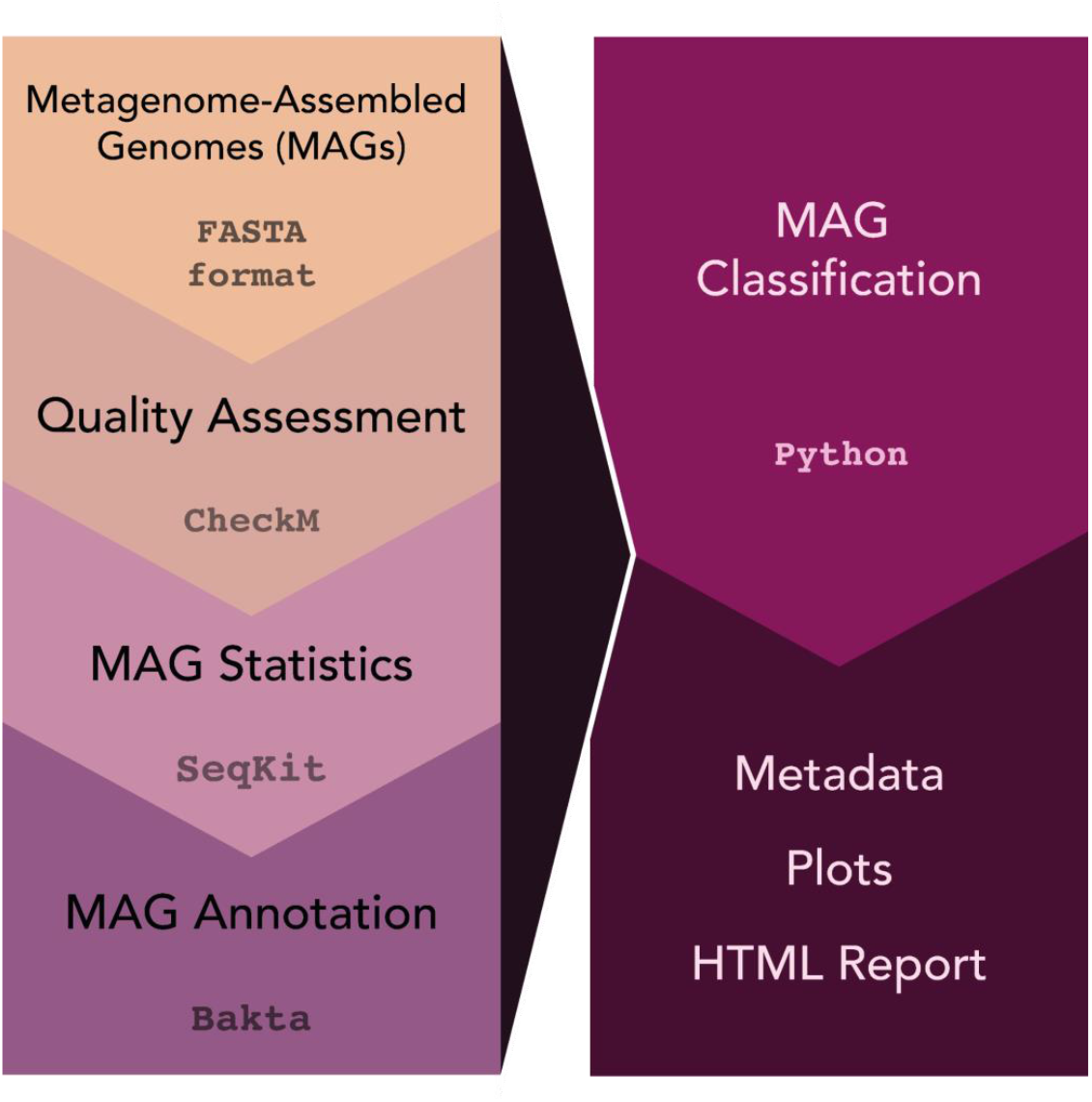
The MAGqual pipeline. MAGs in FASTA format are run through CheckM for quality assessment, SeqKit to calculate basic statistics, and Bakta for MAG annotation, and the output from these is classified according to the MIMAG standards using a bespoke Python script. This outputs a CSV file containing recommended metadata, the MAGs organised according to their classification, and plots and a report are generated to visualise the classification and assembly statistics.

**Figure 2:**
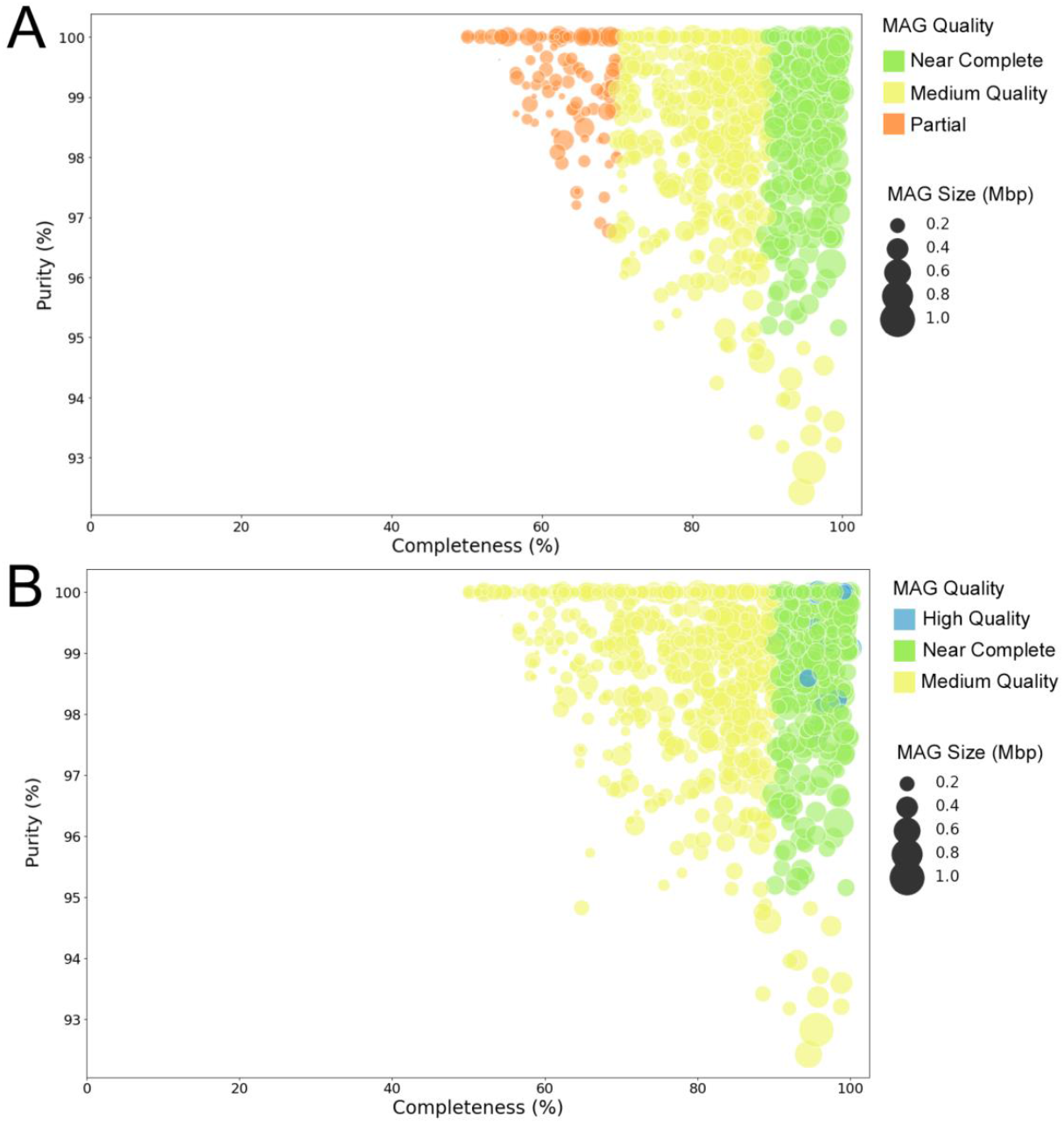
A) The completeness and purity (100-contamination score) of the 1000 MAGs from the metagenomic benchmarking dataset when using the completeness, contamination and quality metrics from the original paper. B) The completeness and purity (100-contamination score) of the 1000 MAGs from the metagenomic benchmarking dataset using the MIMAG standards and the introduced near-complete category

Analysis of this example dataset highlights the importance of the near-complete category in the MAGqual pipeline - all 417 would have been assigned as medium-quality MAGs and only 17 would be classed as high quality, whereas these MAGs are still >90% complete and <5% contaminated and potentially would benefit from further analysis. Including this category in the MAGqual pipeline will hopefully increase the uptake of metagenomic dataset benchmarking in studies. A small number of MAGs have different completeness and purity scores; however, the majority remain the same.

To demonstrate the speed of MAGqual on an HPC, where most metagenomic research is undertaken, these 1,000 MAGs were analysed using the MAGqual pipeline on a 64-core, 512GB machine. MAGqual completed with a wall-clock time of 2:55:57 and a CPU time of 243:34:04.

### Comparison of binning tools

To demonstrate the use of MAGqual as a bin comparison tool a simple gut microbiome metagenome (Shahi *et al*., 2022) was re-binned using three different metagenomic binning tools, CONCOCT, MetaBAT2 and Binsanity (Alneberg *et al*., 2014; Graham, Heidelberg and Tully, 2017; Kang *et al*., 2019) and then refined using the pipeline from MetaWRAP and DAStool (Sieber *et al*., 2018; Uritskiy, DiRuggiero and Taylor, 2018). MAGqual was used to analyse the bins generated using these five different tools. Table 5 shows that CONCOCT and MetaBAT2 both generated a similar number of bins (91 and 92 respectively), however, CONCOCT generated more high-quality bins (8) than MetaBAT2, as did Binsanity. DAStool and MetaWRAP both improved the overall quality of bins, indicating the benefits of a combined binning approach. However, DAStool produced a much lower number of bins overall, and DAStool binned substantially fewer contigs and bases overall (Figure 3A) but produced higher quality MAGs (more complete with low contamination, Figure 3B), illustrating a potential trade-off between assigning more of the sequence data and improved bin quality that likely depends on different algorithmic approaches to binning philosophies. MAGqual enabled a rapid comparison of the MAGs created using these five methods so that users could select the most appropriate binning strategy for their research. Further plots from this analysis can be seen in the HTML file available at https://ac1513.github.io/MAGqual/MAGqual_manuscript_report.html.

**Table 5:** The total number of MAGs and their respective qualities for each of the five binning tools evaluated with this dataset.

|  | Binsanity | CONCOCT | MetaBAT2 | DAStool | MetaWRAP |
| --- | --- | --- | --- | --- | --- |
| Total number of MAGs | 109 | 91 | 92 | 25 | 126 |
| High quality | 8 | 8 | 5 | 10 | 9 |
| Near complete | 1 | 2 | 1 | 2 | 2 |
| Medium quality | 11 | 12 | 18 | 11 | 17 |
| Low quality | 83 | 61 | 66 | 2 | 98 |
| Failed | 6 | 8 | 2 | 0 | 0 |

**Figure 3:**
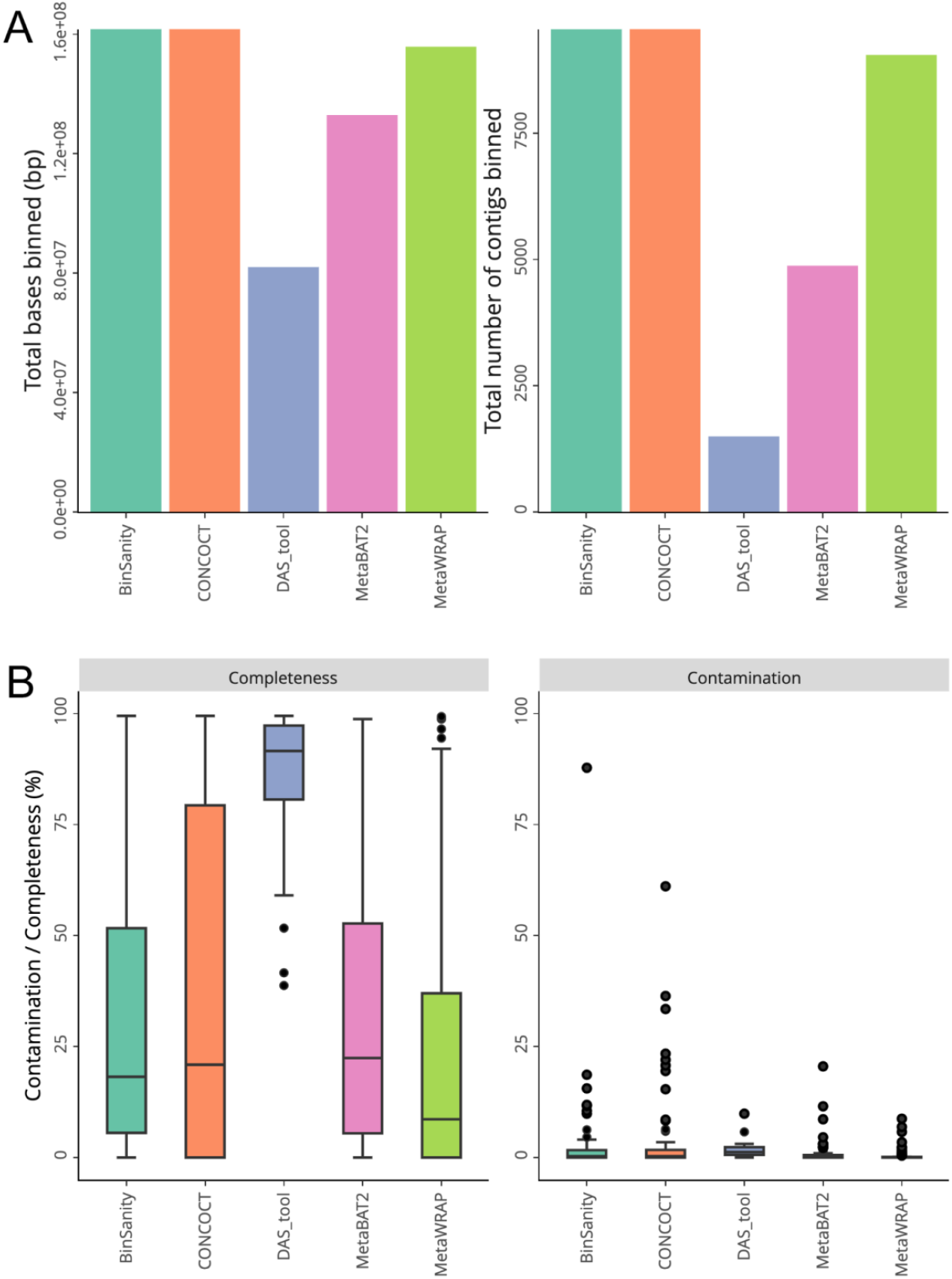
A) The total number of bases (bp) and contigs binned for each of the five binning tools examined with this dataset. B) Boxplot showing the distribution of the completeness and contamination scores for each MAG generated by the five binning tools evaluated with this dataset

## Conclusions

As the size of metagenomic datasets continues to grow, researchers face new challenges in managing and analysing these data. Larger datasets require more sophisticated computational infrastructure, often involving high-performance computing clusters or cloud resources. Metagenome analysis is characterised by a wide array of methods and tools, each with its strengths and limitations, and researchers often choose different tools based on their specific research questions or the nature of their data. This diversity in tools can lead to variations in analysis outcomes. The adoption of MAGqual, a lightweight and user-friendly pipeline, offers a valuable solution to researchers, as it simplifies the binning evaluation process and can be quickly and efficiently applied to datasets of varying sizes generated by any analysis tool.

Building the MAGqual pipeline in Snakemake provides further advantages, including cluster execution, modularisation and simple pipeline updates. For example, during the preparation of this manuscript CheckM2 (Chklovski *et al*., 2023) was released. After testing and benchmarking using the Snakemake benchmarking function, new tools can be integrated into the MAGqual pipeline with ease and, as the pipeline is hosted on GitHub, any updates will be shared with the community promptly.

One of the pipeline’s primary objectives is to encourage wider adoption of the MIMAG (Minimum Information about a Metagenome-Assembled Genome) reporting standards, to ultimately improve the consistency and quality of metagenomic research. MAGqual aids users in swiftly identifying data that merits further analysis. By identifying good quality MAGs, MAGqual provides a targeted approach to metagenome analysis, which can reduce both computational and storage costs, making metagenome analysis more accessible and cost-effective.

## Acknowledgements

We are grateful for computational support from the University of York High-Performance Computing service, Viking and the Research Computing team. JPJC is an Oracle for Research Fellow. We thank Sarah Forrester and Joseph McGrory for their critical reading of the manuscript.

